# FragGeneScanRs: better and faster gene prediction for short reads

**DOI:** 10.1101/2021.08.11.455929

**Authors:** Felix Van der Jeugt, Peter Dawyndt, Bart Mesuere

## Abstract

FragGeneScanRs is a better and faster Rust implementation of the FragGeneScan gene prediction model for short and error-prone reads. Its command line interface is backward compatible and adds extra features for more flexible usage. Compared to the original C implementation, shotgun metagenomic reads are processed up to 22 times faster using a single thread, with better scaling for multithreaded execution.

**Availability and implementation:** The Rust code of FragGeneScanRs is freely available from GitHub under the GPL-3.0 license, with instructions for installation, usage and other documentation.

## Introduction

Studying environmental communities of archaea, bacteria, eukaryotes, and viruses is hampered by problems with isolating and culturing most of these organisms in lab conditions asdf (Locey, K.J. and Lennon, J.T. 2016; Rappé, M.S. and Giovannoni, S.J. 2003; Pedrós-Alió, C. and Manrubia, S. 2016; Hofer, U. 2018; Hahn, M.W., Koll, U., and Schmidt, J. 2019). Metagenomics has therefore become a routine technique to bypass the cultivation step with a combination of high-throughput DNA sequencing and computational methods (Hugenholtz and Tyson 2008; Thomas, T., Gilbert, J., and Meyer, F. 2012). Non-targeted sequencing of genomes in environmental samples, called shotgun metagenomics, in particular allows profiling of both the taxonomic composition and the functional potential of the samples (Quince, C. et al. 2017). Identification of protein coding sequences from shotgun metagenomic reads has therefore become an important precursor to gain insight in the taxonomic and functional diversity of an environmental community (Sharpton, T.J. 2014).

One approach for shotgun metagenomics gene prediction is assembling reads into longer contiguous sequences, called contigs, prior to running traditional gene prediction tools (Breitwieser, F.P., Lu, J., and Salzberg, S.L. 2017). This eases gene prediction, but assembling reads from complex samples with many species in different abundances is a challenging problem. It requires special-purpose algorithms that can be slow, produce artificial contigs, and miss low-abundance genomes (Ghurye, J., Cepeda-Espinoza, V., and Pop, M. 2016; Vollmers, J., Wiegand, S., and Kaster, A. 2017). Direct gene prediction on individual reads can mitigate assembly problems, speed up computations, and enable profiling of low-abundance organisms that cannot be assembled *de novo*. But, it does have to face partial protein coding fragments with missing start/stop codons and read errors (Quince, C. et al. 2017). The choice between assembly-based versus read-based gene prediction may depend on the sample type and the research question at hand.

Various gene prediction tools specialize in directly calling short reads (Hyatt, D. et al. 2012; Hoff, K.J. et al. 2009; Zhu, W., Lomsadze, A., and Borodovsky, M. 2010; Noguchi, H., Taniguchi, T., and Itoh, T. 2008). FragGeneScan (FGS) is the most accurate and popular tool that is currently available (Trimble, W.L. et al. 2012). It uses a hidden Markov model (HMM) that incorporates codon usage bias, start/stop codon patterns, and sequencing error models to predict complete or partial genes in short error-prone reads. Back in 2010, the gene prediction model of FGS was implemented in C and Perl (Rho, M., Tang, H., and Ye, Y. 2010). Multithreading support was added in release 1.19 (August 2014) and most Perl code was replaced with C functions in release 1.30 (April 2016). Because FGS was rather slow, a team of researchers at the University of British Columbia (Canada) forked FGS (presumably from release 1.19) to implement FragGeneScan-Plus (FGS+): a pure C implementation with speedups for single threaded execution and better scaling for multithreading (Kim et al., n.d.).

Both FGS and FGS+ now have pure C implementations that support parallel execution, but their latest releases suffer from their own issues. FGS implements multithreading in a very inefficient way, making it much slower than FGS+. The implementation doesn’t preserve input order, breaking for example the synchronisation between pair-end read files without an extra sorting step in postprocessing. Bugs have also been introduced when replacing Perl code with C functions. In addition, out-of-bound memory access may corrupt its results and cause the software to crash.

FGS+ has inherited inefficient memory usage from FGS and made it worse by copying immutable data structures to individual threads and introducing leaks in the allocated memory. Its multithreading model is overly complex and may deadlock due to race conditions in thread semaphores especially when using a larger number of threads. Although FGS+ is faster than FGS, its results may significantly deviate due to bugs introduced in the reimplementation and missing bug fixes from later FGS versions. For example, FGS+ systematically makes wrong translations of genes encoded on the reverse strand, may result in out-of-bound memory access when copying FASTA headers from a dynamically allocated global array to thread-specific arrays that are statically allocated, uses fixed-length string buffers of 1MB for DNA sequences that overflow for complete genomes, and also crashes when reading from standard input and writing to standard output. In conclusion, FGS generates more accurate results but is slower, whereas FGS+ is considerably faster but generates wrong output.

The source code of FGS and FGS+ is no longer actively maintained, but as metagenomics datasets continue to grow in size, we still need faster gene predictors. In this manuscript we present FragGeneScanRs (FGSrs), a reliable, high-performance, and accurate Rust implementation of the FGS gene prediction model. We ran a benchmark to show that FGSrs produces the same results as FGS and is faster than both FGS and FGS+.

## Implementation

FGSrs is implemented in Rust, a programming language known for its focus on speed and memory-efficiency. In addition, segmentation faults that occur while running FGS or FGS+ are automatically avoided because memory-safety and thread-safety are guaranteed by Rust’s type system and ownership model. Its zero-cost abstractions yield more readable code and put optimizations in the hands of the compiler.

We started off with a Rust implementation that was equivalent to FGS release 1.31. Afterwards, we gradually optimized performance and improved the quality of the software, while monitoring equivalence with the original implementation. We outline some of these optimizations and improvements in what follows and refer to the source code and its documentation for more details.

FGS uses statically allocated 300 KB buffers for different representations of reads and 100 KB buffers for protein translations. By using dynamically allocated buffers that grow when needed instead, we avoid static buffer overflows, increase speed, and reduce the memory footprint.

FGS stores default training data for its HMM in separate text files. We include a binary representation of default data in the executable during compilation, but still support passing custom training data as command line arguments. This improves usability by reducing dependencies during installation of the software and allowing to execute FGSrs with default training data anywhere on the system without the need to run FGS in its own directory or explicitly pass the path to the default training files (new option -`r`).

The HMM configurations used by FGS are immutable after initialization, but the original implementation wastes memory by copying them to each thread. We store this data in shared memory that threads can access concurrently. We use mutexes for protected access to shared input and output file handles. This avoids the need to split input in chunks upfront, have threads that store results per chunk in a separate file, and merge these files afterwards. This eliminates disk overhead and speeds up I/O.

HMM gene regions have six inhomogeneous sets of states that represent matches, insertions and deletions for two successive codons in a read. FGS processes the six states in a loop, combined with conditional execution to handle topological differences between state transitions. Unrolling these loops not only makes the condition-less code more readable, it is also significantly faster. We have further improved readability of the code by replacing #define constants in C with Rust enum types.

Where FGS uses row-major order to store dynamic programming matrices, we switched to column-major order for improved locality when accessing matrix elements in the Viterbi algorithm. This results in a speedup because the reordered memory layout causes less cache misses.

FGS outputs the DNA sequence, its translated amino acid sequence, and additional metadata for each coding region found in a read, with optional formatting to indicate insertions and deletions in the DNA sequence. This requires FGS to compute and store the DNA sequence twice during the backtracking step at the end of the Viterbi algorithms. Once for reconstructing the formatted DNA sequence and once for reconstructing the unformatted DNA sequence. We produce a unified representation and delay formatting until output is generated. We also note that a bug was introduced in FGS (release 1.30) when the backtracking step was converted from Perl to C, which generates DNA and protein sequences for complete genomes that are incorrect.

## Results

The FGSrs command line interface is backward compatible with FGS, so it can be used as a faster and memory-friendly drop-in replacement for FGS in bioinformatics pipelines. FGSrs also has some additional features that enable more flexible usage. We support the conventional standard input and output channels as alternatives for passing files as arguments to the -`s` and -`o` options. This enables embedding FGSrs in POSIX pipes without storing intermediate results on the file system. Storage locations for generated DNA sequences (option -`n`), translated amino acid sequences (option -`a`) and metadata (option -`m`) can also be specified individually, which may give additional speedups because unspecified information does not need to be computed. Where FGS and FGS+ report gene predictions out-of-order, FGSrs by default preserves read order. The priority queue that guarantees in-order reporting comes with a small overhead on speed and memory usage, but can be disabled with the option -`u`.

We ran a benchmark on a 16-core Intel® Xeon® CPU E5-2650 v2 at 2.60 GHz, to evaluate the performance improvements of FGSrs (release 1.0.0) over FGS (release 1.31) and FGS+ (git version 91b0ab6). The sample datasets included in the FGS and FGSrs repositories are used as input data and measurements are averaged over 5 runs. When all three implementations are executed single threaded (Figure 1), FGSrs processes short reads (80 bp) 22.6× faster than FGS and 1.2× faster than FGS+. Long reads (1,328 bp) are processed 4.2× times faster than FGS and 1.6× faster than FGS+. The bulk of the runtime is consumed by the Viterbi algorithm, having a time complexity of *O*(*s*^2^ × *n*) with *s* the number of HMM states and *n* the length of the sequence that needs to be processed. Because the HMM of the gene prediction model has a fixed number of states (*s* = 49), we can thus expect runtime to grow linearly for reads that are even longer. FGSrs processes the complete genome sequence of *Escherichia coli* str. K-12 subst. MG1665 (NC_000913; 4,639,675 bp) 2.2× faster than FGS and 347.6× faster than FGS+. The latter measurement essentially shows that FGS+ is not fit for processing complete genomes in practice. FGSrs and FGS+ scale better for multithreaded execution than FGS (Figure 2). Increasing the thread count from 1 to 8, this results in a speedup factor of 6.5 for both FGSrs and FGS+ and only 5.4 for FGS. The difference increases for higher thread counts and the execution even consistently halts due to race conditions for FGS+ when using more than 10 threads. FGSrs and FGS have a comparable memory footprint of about (80 + 70*t*) MB for *t* threads (Figure 3). FGS+ consumes more memory with about (265 + 85*t*) MB for *t* threads. However, the memory requirements are not a limiting factor to run any of these tools on a standard laptop, with 4 threads needing between 350MB and 520MB RAM.

**Figure 1:**
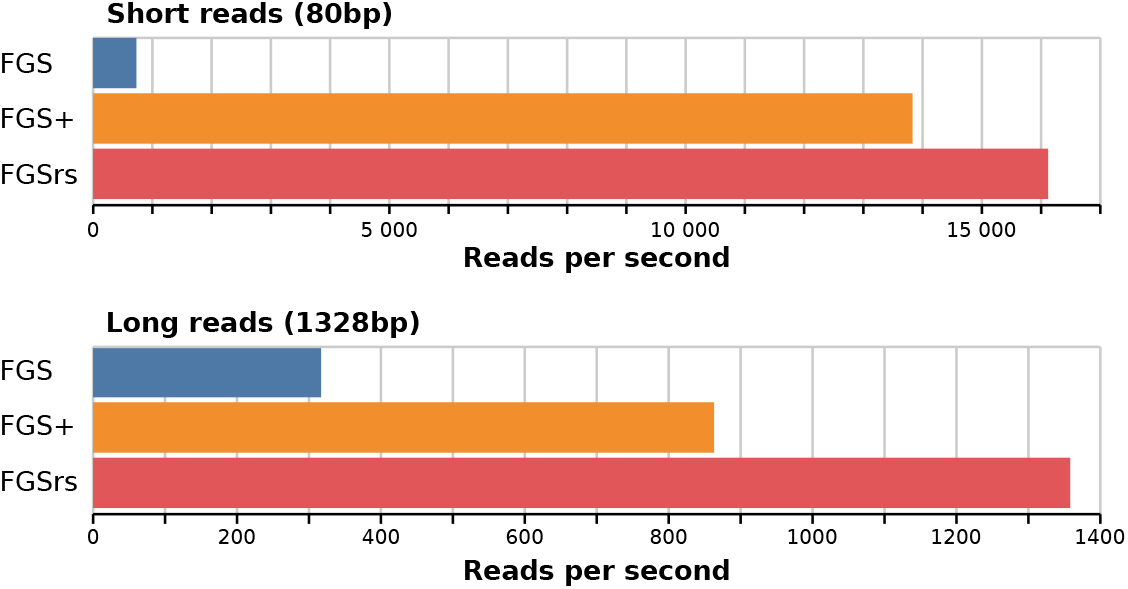
Processing speed for single threaded execution of FGS, FGS+ and FGSrs on short (80 bp) and long (1,328 bp) reads. FGS and FGSrs generate DNA sequences, protein translations and metadata, whereas FGS+ only generates protein translations because the software crashes when other output is generated. FGS and FGS+ report gene predictions out-of-order, where default in-order reporting was used for FGSrs.

**Figure 2:**
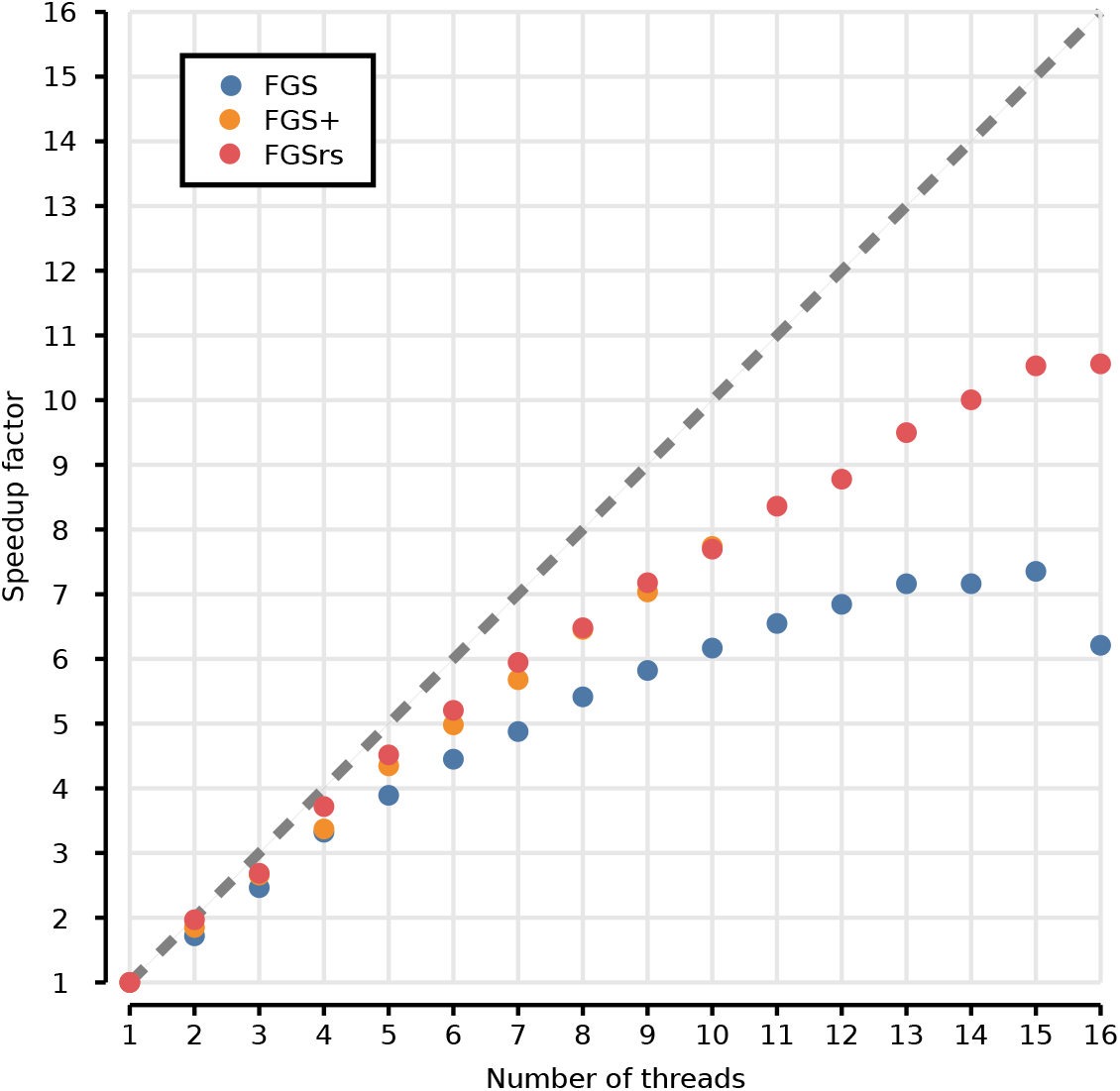
Scaling for multithreaded execution of FGS, FGS+ and FGSrs on long reads (1,328 bp), computed as the speedup of concurrent execution with *t* threads (*x*-axis) over single threaded execution. Dash line shows the theoretical upper bound for the speedup. Race conditions consistently halt the execution of FGS+ above 10 threads. FGS and FGSrs generate DNA sequences, protein translations and metadata, whereas FGS+ only generates protein translations because the software crashes when other output is generated. FGS and FGS+ report gene predictions out-of-order, where default in-order reporting was used for FGSrs.

**Figure 3:**
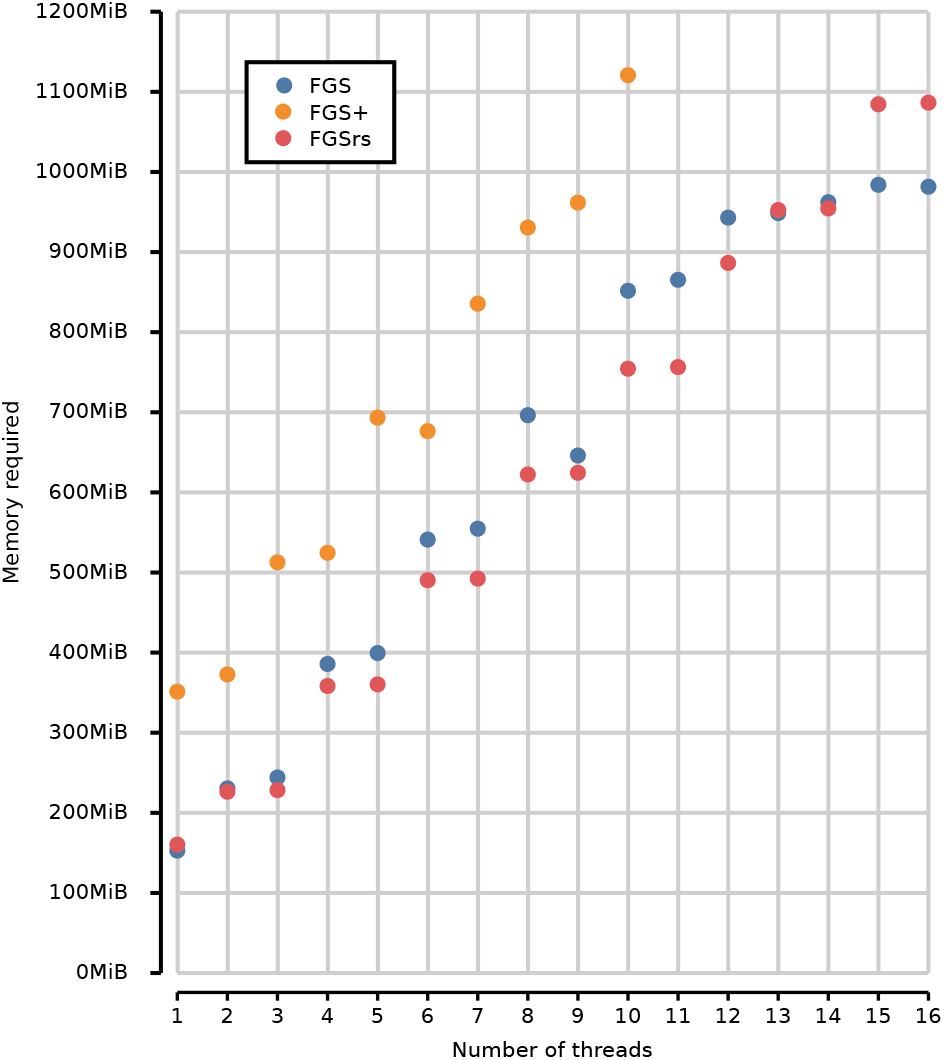
Memory footprint for multithreaded execution of FGS, FGS+ and FGSrs on long reads (1,328 bp). Total memory footprint (heap, stack and memory-mapped file I/O) measured using the Massif heap profiler of Valgrind (Nethercote, N. and Seward, J. 2007) with --`pages-as-heap` option. Race conditions consistently halt the execution of FGS+ above 10 threads. FGS and FGSrs generate DNA sequences, protein translations and metadata, whereas FGS+ only generates protein translations because the software crashes when other output is generated. FGS and FGS+ report gene predictions out-of-order, where default in-order reporting was used for FGSrs.

In conclusion we can state that FGSrs is a reliable implementation of the FGS gene prediction model that is an order of magnitude faster than the original implementation. Its command line interface is backward compatible with extensions for more flexible usage. The source code of FGSrs is freely available from GitHub under the GPL-3.0 license, with instructions for installation, usage and other documentation.

## Acknowledgements

This work was supported by the Research Foundation–Flanders (FWO) (12I5220N to B.M.) We thank the students of the Computational Biology class of 2019-2020 for scrutinizing issues with the code of FGS and FGS+.

## References

Breitwieser, F.P., Lu, J., and Salzberg, S.L. 2017. “A Review of Methods and Databases for Metagenomic Classification and Assembly.” Briefings in Bioinformatics 5: 1125–36. https://doi.org/10.1093/bib/bbx120.

Ghurye, J., Cepeda-Espinoza, V., and Pop, M. 2016. “Metagenomic Assembly: Overview, Challenges and Applications.” The Yale Journal of Biology and Medicine 20: 353–62.

Hahn, M.W., Koll, U., and Schmidt, J. 2019. The Structure and Function of Aquatic Microbial Communities. Cham: Springer International Publishing. https://doi.org/10.1007/978-3-030-16775-2_10.

Hofer, U. 2018. “The Majority Is Uncultured.” Nature Reviews Microbiology 16: 716–17.

Hoff, K.J., Lingner, T., Meinicke, P., and Tech, M. 2009. “Orphelia: Predicting Genes in Metagenomic Sequencing Reads.” Nucleic Acids Research 28: W101–5. https://doi.org/10.1093/nar/gkp327.

Hugenholtz, P., and G. W. Tyson. 2008. “Metagenomics.” Nature 455: 481–83.

Hyatt, D., LoCascio, P.F., Hauser, L.J., and Uberbacher, E.C. 2012. “Gene and Translation Initiation Site Prediction in Metagenomic Sequences.” Bioinformatics 12: 2223–30. https://doi.org/10.1093/bioinformatics/bts429.

Kim, D.J., Hahn, A.S., Wu, S.J., Hanson, N.W., Konwar, K.M., and Hallam, S.J. n.d. “FragGeneScan-Plus for Scalable High-Throughput Short-Read Open Reading Frame Prediction.” In.

Locey, K.J., and Lennon, J.T. 2016. “Scaling Laws Predict Global Microbial Diversity.” Proceedings of the National Academy of Sciences 113: 5970–75. https://doi.org/10.1073/pnas.1521291113.

Nethercote, N., and Seward, J. 2007. “Valgrind: A Framework for Heavyweight Dynamic Binary Instrumentation.” In. https://doi.org/10.1145/1250734.1250746.

Noguchi, H., Taniguchi, T., and Itoh, T. 2008. “MetaGeneAnnotator: Detecting Species-Specific Patterns of Ribosomal Binding Site for Precise Gene Prediction in Anonymous Prokaryotic and Phage Genomes.” DNA Research 38 (6): 387–96. https://doi.org/10.1093/dnares/dsn027.

Pedrós-Alió, C., and Manrubia, S. 2016. “The Vast Unknown Microbial Biosphere.” Proceedings of the National Academy of Sciences 113: 6585–87. https://doi.org/10.1073/pnas.1606105113.

Quince, C., Walker, A.W., Simpson, J.T., Loman, N.J., and Nicola, S. 2017. “Shotgun Metagenomics, from Sampling to Analysis.” Nature Biotechnology 35: 833–44. https://doi.org/10.1038/nbt.3935.

Rappé, M.S., and Giovannoni, S.J. 2003. “The Uncultured Microbial Majority.” Annual Review of Microbiology 57: 369394. https://doi.org/10.1146/annurev.micro.57.030502.090759.

Rho, M., Tang, H., and Ye, Y. 2010. “FragGeneScan: Predicting Genes in Short and Error-Prone Reads.” Nucleic Acids Res. 38 (20): e191.

Sharpton, T.J. 2014. “An Introduction to the Analysis of Shotgun Metagenomic Data.” Frontiers in Plant Science, 209. https://doi.org/10.3389/fpls.2014.00209.

Thomas, T., Gilbert, J., and Meyer, F. 2012. “Metagenomics - A Guide From Sampling to Data Analysis.” Microbial Informatics and Experimentation 2. https://doi.org/10.1186/2042-5783-2-3.

Trimble, W.L., Keegan, K.P., D’Souza, M., Wilke, A., Wilkening, J., Gilbert, J., and Meyer, F. 2012. “Short Read Reading-Frame Predictors Are Not Created Equal: Sequence Error Causes Loss of Signal.” BMC Bioinformatics 15 (1). https://doi.org/10.1186/1471-2105-13-183.

Vollmers, J., Wiegand, S., and Kaster, A. 2017. “Comparing and Evaluating Metagenome Assembly Tools from a Microbiologist’s Perspective - Not Only Size Matters!” PLOS ONE 89: 131. https://doi.org/10.1371/journal.pone.0169662.

Zhu, W., Lomsadze, A., and Borodovsky, M. 2010. “Ab Initio Gene Identification in Metagenomic Sequences.” Nucleic Acids Research 37 (12): e132–32. https://doi.org/10.1093/nar/gkq275.

